# Outer Hair Cell Function is Normal in βV Spectrin Knockout Mice

**DOI:** 10.1101/2021.02.01.427491

**Authors:** Michael C. Stankewich, Jun-Ping Bai, Paul R. Stabach, Saaim Khan, Winston J.T. Tan, Alexei Surguchev, Lei Song, Jon S. Morrow, Joseph Santos-Sacchi, Dhasakumar S. Navaratnam

## Abstract

Reports have proposed a putative role for βV spectrin in outer hair cells (OHCs) of the cochlea. In an ongoing investigation of the role of the cytoskeleton in electromotility, we tested mice with a targeted exon deletion of βV spectrin (Spnb5), and unexpectedly find that Spnb5^(-/-)^ animals’ auditory thresholds are unaffected. Similarly, these mice have normal OHC electromechanical activity (otoacoustic emissions) and non-linear capacitance. In contrast, magnitudes of auditory brainstem response (ABR) wave 1-amplitudes are significantly reduced. Evidence of a synaptopathy was absent with normal hair cell CtBP2 counts. In Spnb5^(-/-)^ mice, the number of afferent and efferent nerve fibers is decreased. Consistent with this data, Spnb5 mRNA is present in Type I and II spiral ganglion neurons, but undetectable in OHCs. Together, these data establish that βV spectrin is important for hearing, affecting neuronal structure and function. Significantly, these data exclude βV spectrin as functionally important to OHCs as has been previously suggested.

## INTRODUCTION

In the early 1990s, several groups made key seminal observations of a spectrin/actin lattice in the cochlear outer hair cells (OHCs) (Holley & Ashmore, 1990) (Holley, Kalinec, & Kachar, 1992). Their electron micrographs propelled an exquisite model of the spectrin/actin lattice assembled just beneath the OHC lateral membrane with longitudinal spectrin molecules linked to circumferential actin filaments that, in turn, were linked to the plasma membrane by electron-dense pillars. Their work also identified a pool of spectrin in the actin-rich cuticular plate. Another study evaluated spectrin proteins in cochlear innervation during rodent development (Hafidi, Despres, & Romand, 1990). They identified differences in developmental expression of the erythroid beta spectrin (Spnb1 spectrin) compared to the most ubiquitous nonerythroid form αII/ βII spectrin, similar to what is observed in the brain. What is not certain from these earlier studies is the specific spectrins involved, a family now known to consist of two alpha spectrins (Spna1 and 2) and five beta spectrins (Spnb1-5). More recently, it has been shown that Spnb4 is restricted to the nodes of Ranvier in myelinated axons and is also a deafness-related gene (Parkinson et al., 2001) (Lewis et al., 2018). It appears from these studies that the deafness in the quivering mouse is likely not originating in the cochlea but rather is central in origin. Furthermore, Spnb2 spectrin, when selectively deleted from the mouse cochlea sensory epithelia by crossing Spnb2 floxed mice with Atoh-1 Cre-recombinase, causes profound deafness due to disruption of the stereocilia rootlets in the cuticular plate in hair cells (Liu et al., 2019). Thus, the function of specific spectrins, particularly as it relates to the OHC lateral membrane and cochlear innervation, remains unclear. In humans, the molecule βV spectrin (SPTBN5) that was first discovered in the retina, is involved with the retinal disorder, Usher syndrome (Stabach & Morrow, 2000) (Papal et al., 2013). Further reports have proposed a putative role for βV spectrin in a second sensorineural function, hearing. One group’s observations demonstrated that antisera raised against the C-terminal peptide of SPTBN5, labeled structures along the lateral membrane of the OHC (Legendre, Safieddine, Kussel-Andermann, Petit, & El-Amraoui, 2008) (Cortese et al., 2017). This observation drew attention because detection of this giant spectrin in tissue other than retina and testis has been elusive. This finding raised the possibility that βV spectrin was involved in the electromotile response of OHCs.

Continuing an ongoing effort in our lab to define the cytoskeleton’s role in electromotility (Bai et al., 2010), we sought to determine the role of βV spectrin in OHC function. In this study, we analyzed βV spectrin expression in the cochlea and addressed its function in mice with a targeted deletion of Spnb5. We tested hearing by auditory brainstem responses (ABRs), measured OHC-mediated otoacoustic emissions and OHC nonlinear capacitance (NLC), and evaluated the number of inner hair cell (IHC) ribbon synapses and nerve fibers that innervate the cochlear sensory epithelium. We find that while OHC function is unaffected in the absence of βV spectrin, nerve counts are reduced at the habenular, in all three turns of the cochlea, and in efferent fibers in the mid and basal turns, which collectively account for the reduced wave I amplitudes in ABRs. Using RNAscope *in situ* hybridization, we confirm expression of Spnb5 mRNA in Type I and II SGNs and absent expression in OHCs. Moreover, mRNA transcript was significantly reduced in the SGN of Spnb5^(-/-)^ mice, confirming knockdown of the gene. We conclude that βV spectrin is necessary to maintain the integrity or function of afferent and efferent nerves of the peripheral auditory system, but plays no role in the OHC itself.

## RESULTS

### Mouse βV spectrin amino acid sequence is divergent from the human protein

Surprisingly, a sequence alignment comparing the mouse amino acid sequence to the human amino acid sequence reveals a significant divergence. In contrast, other beta spectrins, including beta I-IV, have ~ 90% amino acid conservation between human and mouse sequences. By aligning human and mouse βV spectrin, we note a significant divergence at the N- and C-termini of the proteins and an internal stretch of amino acids from position 817-922 (**Figure 1A**). Of note is that the initiator methionine residue in the human form is conserved between mouse and human genes and lies at position 36 and within exon 2 (**Figure 2**). The mouse sequence incorporates an unusual homorepeat of glutamines and, notably, does not have a bona fide pleckstrin homology (PH) domain seen in the human form.

**Figure 1.**
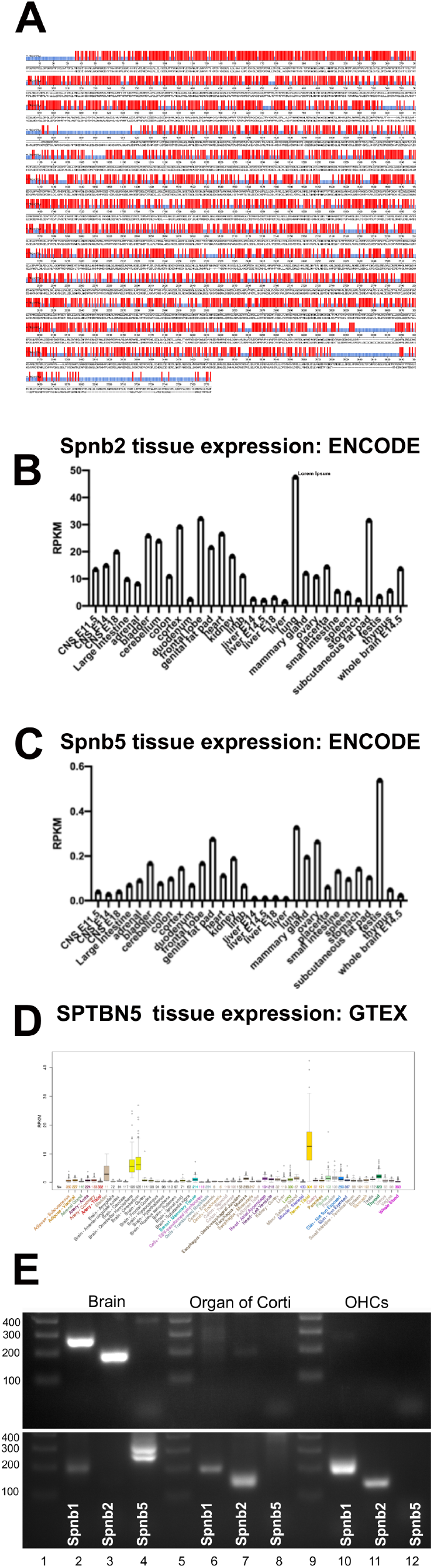
**(A) Spnb5 mouse** (<u>ENSMUST00000156159.4</u>) **amino acid sequence differs from humans.** Shown is the comparison of the deduced amino acid sequence of mouse (lower) and human (upper) Spnb5. Conserved amino acids are marked in red. There is divergence at the N and C termini. In addition, a region in the middle of the protein from posiiton 817-922 shows significant divergence. The mouse version of this gene shares only 65.6 percent identity with the human form. **(B, C) Spnb5 is expressed in brain and testis.** The relative abundance of Spnb2 transcripts as compared to Spnb5 in mouse tissues. Shown are the expression of Spnb2 and 5 in RNA Seq. NCBI reference datasets (ENCODE; Bioproject PRJNA66167_(Yue et al., 2014). On average, the abundance of Spnb5 mRNA is an order of magnitude less abundant than Spnb2. Of the tissues examined, adult testis has the most mRNA for Spnb5. **(D) In human, Spnb5 is most expressed in peripheral nerve.** The relative abundence of Spnb5 in different tissues in GTEX database is presented (version 6) (Consortium, 2013). Of all the tissues examined Spnb5 is expressed in highest amounts in the peripheral nerve (tibial nerve). **(E) Spnb5 mRNA is not detectable in the organ of Corti or OHCs.** Shown are PCR amplifications from mouse brain (lanes 2-4), organ of Corti (lanes 6-8) and OHCs (lanes 10-12) of Spnb1 (lanes 2, 6 and 10), Spnb2 (lanes 3, 7 and 11) and Spnb5 (lanes 4, 8 and 12). The upper panel shows amplicons for these three mRNA transcripts using outer primers while the lower panel shows amplicons of primers using a nested set of primers. All three spectrins are detectable in brain. Spnb1 and 2 are detectable in both the organ of Corti and OHCs (lanes 6-7 and 10-11). Spnb5 however, is not detectable using

**Figure 2.**
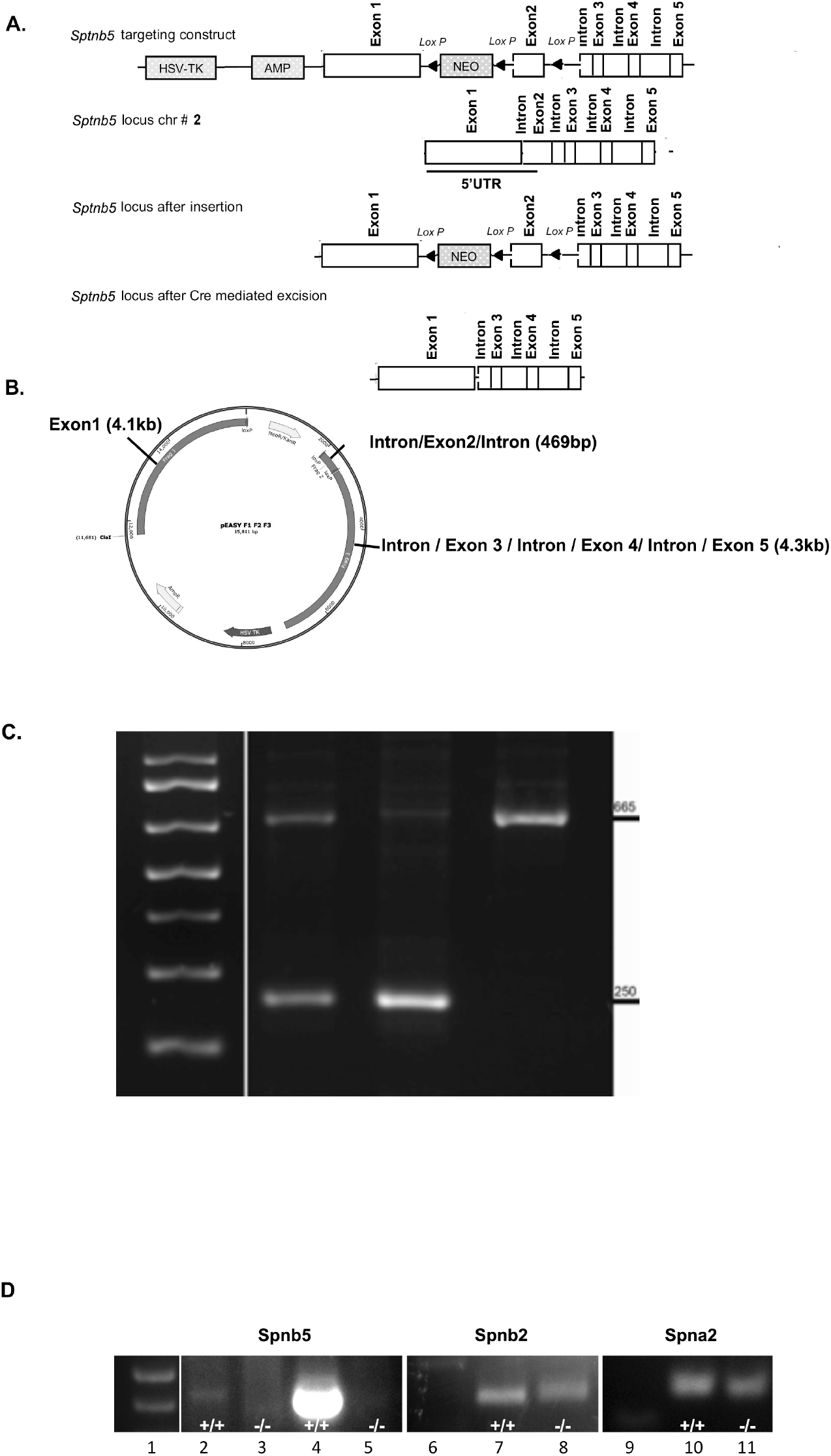
Exon 2 of the Spnb5 gene was removed using homologous recombination. **(A) Schematic of the strategy used to delete exon 2.** The construct contained a short arm of exon 1 and a long arm of exons 3, 4, and 5 together with the intervening introns. Both the selection marker Neo and exon two are flanked by Lox P sites. Linearized plasmid was injected into ES cells and clones with homologous recombination and selected by neomycin (Neo)/ G418 resistance. Mice were then crossed with Actin-Cre mice to generate floxed mice lacking the Neo cassette and exon 2. **(B) Vector map of pEASY used to target Spnb5 deletion.** Exon 2 was subcloned between two Lox P sites adjacent to the Neo gene. These two genomic fragments were in turn placed between a 4.1kb short arm containing Exon 1 and a 4.3kb long arm containing exons 3, 4, and 5 and the intervening introns. The vector also contains a thymidine kinase and ampicillin gene allowing for selection. The plasmid was linearized with Cla1. **(C) PCR amplification of genomic DNA confirming deletion of exon two and flanking intronic sequence.** In genomic DNA from Spnb5^(-/-)^ mice, primers that amplify a product no longer contain part of intron 1; exon 2; and part of intron 2/3. Lane 1 base pair standards, lane 2 Spnb5^(+/-)^, lane 3 Spnb5^(-/-)^, lane 4 Spnb5^(+/+)^. The larger fragment of 665 bp consists of unexcised exon two while the smaller fragment consists of genomic DNA fragment from which exon two has been excised. **(D) Decreased expression of Spnb5 in Spnb5^(-/-)^ testicular cDNA compared to Spnb2 and Spna2**. Spnb5 exons 58-60 in lanes 2-3, Spnb5 exon 53-55 in lanes 4-5; Spnb2 in lanes 7-8 (exons 28-29) and Spna2 (exons 54-55) in lanes 10-11. WT in lanes 2, 4, 7 and 10. Spnb5^(-/-)^ in lanes 3, 5, 8 and 11.

### Spnb5 mRNA tissue expression

Spnb5 spectrin is expressed in most mouse tissues but at low levels. In ENCODE, the level of Spnb5 transcripts is an order of magnitude less abundant than the ubiquitous form Spnb2 (**Figure 1B**) (Yue et al., 2014). The peak abundance of Spnb2 approaches 50 RPKM while Spnb5 is 0.5 RPKM in testis (**Figure 1B, C**). Interestingly, the GTEX dataset shows high expression of Spnb5 in the peripheral (tibial) nerve (**Figure 1D**). To validate that mRNA is detected in mice, we tested RNA from the testis with primer sets that bridged exons. By standard RT-PCR, Spnb5 mRNA fragments encoding both the N-terminus and C-terminus were amplified from testis and brain RNA, and its identity was confirmed by Sanger sequencing. In contrast to brain and testicular tissue, we could not detect Spnb5 mRNA in isolated OHCs and whole organ of Corti by both conventional and nested PCR (Table 1 details the gene-specific primer sets). Using RT-PCR from ~3000 OHCs and organ of Corti, we detect both Spnb1 and Spnb2. Using nested PCR, amplicons for these spectrins are robust, while we do not visualize any product for Spnb5 (**Figure 1E**). These data are in accord with data derived from the public data sets of transcriptome profiling for OHCs in gEAR (https://umgear.org/) (**See supplementary Figure1**). Furthermore, *in situ* hybridization for Spnb5 using RNAscope corroborates these findings. We are unable to detect Spnb5 mRNA in OHCs (**see Supplementary Figure 2, panels C and S**), whereas mRNA for the potassium channel, Kcnma1, is readily detected in OHCs (green signal in **panel B**). In contrast, the Spnb5 probe is detected in spiral ganglion neurons (SGNs) (red signal in both **panels C and P**). To further characterize the expression of Spnb5 in SGNs, we co-labeled Spnb5 (green) with known markers of myelinated Type I ganglion cells (tyrosine hydroxylase, (TH)) and unmyelinated Type II ganglion cells (Epha4) in red (see Supplementary Figure 3). We find that Spnb5 signal overlaps with the signals from both TH (panel A) and Epha4 (panel B). These data demonstrate that Spnb5 is expressed in both Type I and II SGNs.

**Table1.**
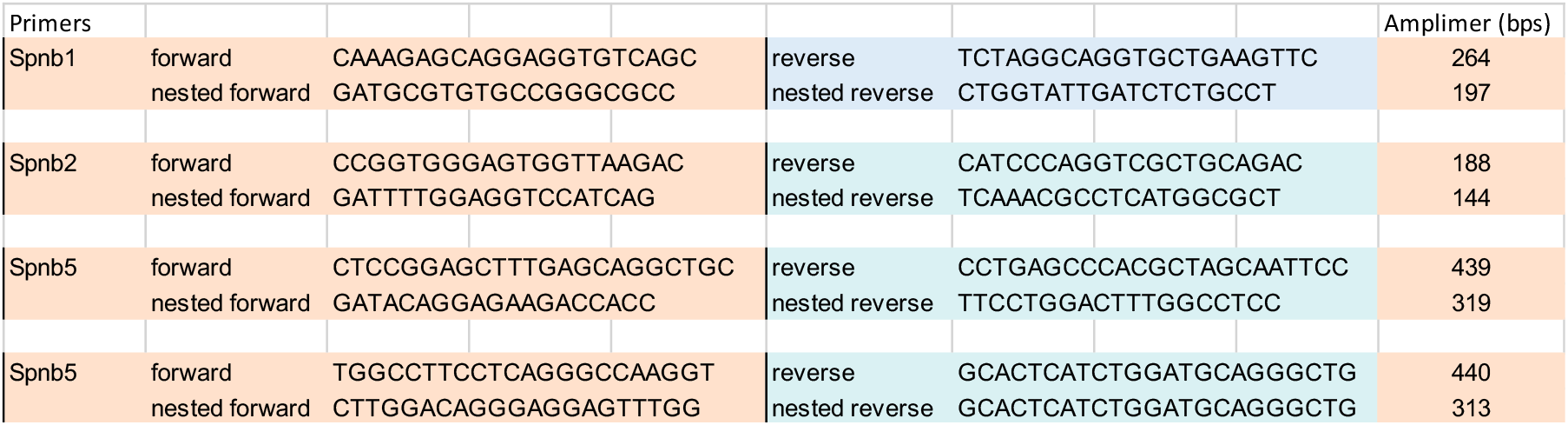
Beta spectrin gene specific primers. Primer sets were used to detect for mRNA of Spnb1, Spnb2, and Spnb5 in the brain, organ of Corti, and isolated OHCs. The PCR amplimer sizes from mRNA are presented in the final column.

**Figure S1.**
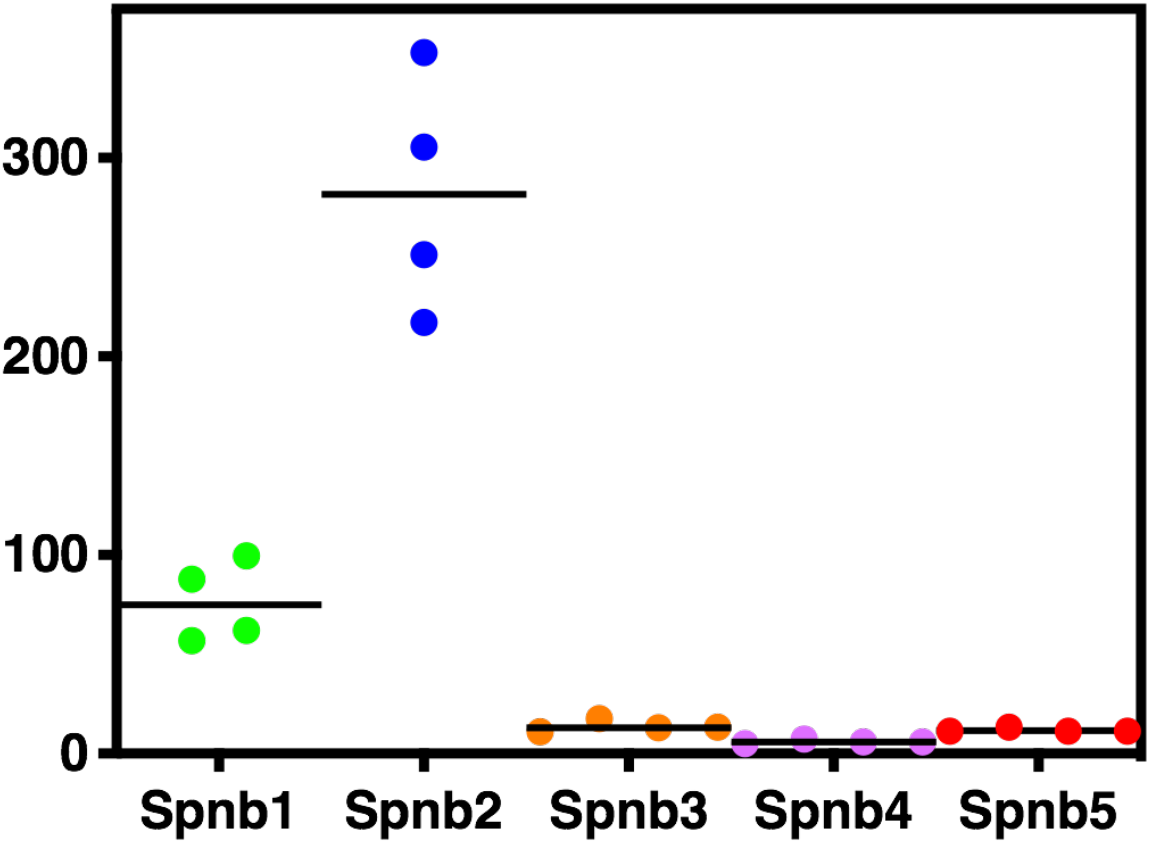
Expression levels for each of the beta spectrins in OHCs. Mining data from the gene Expression Analysis Resource (gEAR) we graphed the abundance of each of the five beta spectrin genes. This data set was derived from transcriptome analysis using sc-RNAseq of cell populations isolated from mouse cochleae (Li et al., 2018).

### Targeted knockout of Spnb5 in mice using homologous recombination

In order to properly target the βV spectrin gene in mice, we needed to know its precise gene structure. While the human gene has been well characterized and indexed since its discovery in 2000 (Stabach & Morrow, 2000), there has been ambiguity in the intron/exon usage in the mouse gene found on chromosome 2. The GENBANK entry for Spnb5 has been modified many times in the last several years, with the last significant annotation in 2007 (Benson et al., 2013). This annotation was derived by automated computational analysis using gene prediction algorithms. The exon prediction is made difficult because 1) the gene is very large, so PCR amplifying a full-length cDNA is not trivial, and 2) the conservation between human and mouse intron/exon boundaries is weak at the N- and C-termini. The current and accepted annotated version in both the UCSC genome browser and in Ensembl shows Spnb5 spectrin mRNA in the mouse to be 14.4 kb with a 10,872 bp ORF that has 66 exons and codes for 3623 amino acids: NM_001370938: <u>NP 001357867.1.</u> This version has been validated in the ENCODE project based on the robustness of next-generation sequencing efforts over the last several years (Yue et al., 2014).

Shown in the schematic in Figure 2A is the strategy for targeted deletion of exon 2 of Spnb5. The knockout strategy was based on the transcript ID: ENSMUST00000156159.3, and a targeting construct was generated in which three Lox P sites were inserted to flank the neomycin selection marker and exon 2 containing the start codon, resulting in disruption of the open reading frame upon Cre-mediated excision. Exon 2 contains the ATG start codon (with a consensus Kozak sequence) and an additional 37 amino acids at its N-terminus. Notably, the codon for the methionine residue corresponding to the human start codon is also contained within the second exon. A genomic bacterial artificial chromosome containing the Spnb5 locus was obtained, and genomic DNA fragments containing the first exons of Spnb5 were subcloned in pEasy (a gift from Sankar Ghosh). The targeting vector comprised a 4.3 kb long arm, a 4.1 kb short arm, and a middle selection cassette which contains exon two flanked by a PGK–neomycin resistance (Neo), two FRT sites, and the herpes simplex thymidine kinase gene (Figure 2B and Primer Sets in Table 2). The targeting plasmid DNA was linearized with Cla1 and electroporated into mouse embryonic stem cells (ESCs) (129SV/EV). ESCs were selected for neomycin/G418 resistance. Positive clones were identified by PCR and confirmed by Southern blot analysis using external and internal probes. A correctly targeted ESC clone was injected into C57BL/6J blastocysts to produce germline transmitting chimeric mice. These were then mated with actin-Cre C57BL/6J transgenic mice for generation of heterozygous Spnb5 transgenic mice with both the Neo cassette and exon two removed (Spnb5^(+/-)^). The resulting Spnb5^(+/-)^ offspring were crossed to generate βV spectrin-deficient mice. The genotypes of these mice were confirmed by PCR amplification (Figure 2C). Chimeric mice were backcrossed onto the C57BL/6J strain. Litters from heterozygous breeders Spnb5^(+/-)^ / Spnb5^(+/-)^ were of normal number, and the mice of all genotypes were phenotypically indistinguishable with normal lifespans >2 yrs. old. We confirmed decreased expression of Spnb5 in cDNA from testes of Spnb5^(-/-)^ mice, where we found equivalent expression of Spnb2 and Spna2 (Figure 2D and Primer Sets in Table 3). The decreased level of Spnb5 mRNA was corroborated by RNAScope in situ hybridization (ISH) experiments. In low power images of cochlear tissue (Supplementary Figure 2, panels A-I), the signal for Spnb5 (red) in the SGN is significantly reduced in KO mice. Quantitation of the mean fluorescence intensity shown in panel V reveals that the amount of signal in SGNs is significantly reduced by greater than 60% in the SGN of Spnb5^(-/-)^ animals. Higher power magnification ISH images, when labeled with Spnb5 probe alone, reveals a significant reduction in both the number of fluorescent particles per neuron in KO tissue (panel W). On average, the number of particles per neuron with positive granules was significantly reduced by greater than ~50%. In addition, both the total fluorescence intensity and the number of neurons that contained Spnb5 is dramatically reduced in the SGNs of Spnb5^(-/-)^ animals (panel C vs. F). The binding site for the RNAscope *in situ* probes (at position 7163 – 8000 in XM_006500537; 3669-4506 in ENSMUST00000156159.4) corresponds to ~ 1300 amino acids after the initiator methionine. These data suggest that the transcript up to position 3669-4506 was significantly reduced in the SGN of knockout animals. It should be noted that the actin binding of spectrins occurs through calponin homology (CH1 and CH2) sites present at the N-terminus of the proteins. In Spnb5, these sites are located at aa 20-124 (CH1) and aa 143-247 (CH2) and encoded by the transcript well before the RNAscope *in situ* probe binding sites. In other beta spectrins, loss of function mutations occur within the CH1 and CH2 actin binding domains (Cousin et al., 2020). While it is still possible that there are mRNA transcripts that are shorter with in-frame methionine start sites, these transcripts would not be expected to produce a functional protein in lieu of it lacking the CH1 and CH2 actin binding domains (there are, for instance, 43 in-frame methionine codons, 19 of which also contain consensus Kozak sequences, that lie downstream of the *in situ* hybridization probe binding sites).

**Table 2.**
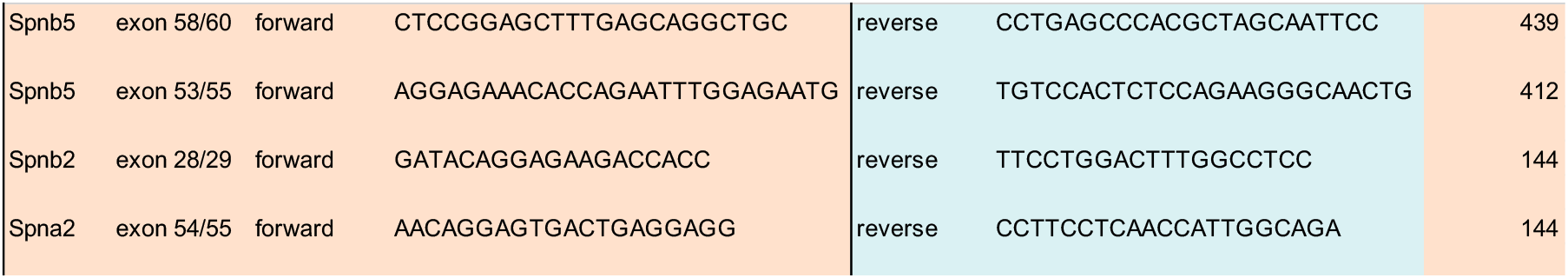
Primers for the Spnb5 gene used to prepare the three DNA fragments that were subcloned into pEasy-Flox. The targeting vector for homologous recombination was constructed by cloning three PCR generated fragments into the pEASY-Flox vector. Fragment 1 is a 4115 bp product, fragment 2 is a 469 bp product that contains Spnb5 exon 2, and fragment 3 is 4371 bp of sequence that spans up to exon 5. The three fragments were sequence verified. The oligos contain common restriction sites at their 5’ end for easy cloning into the targeting vector.

**Table 3.**
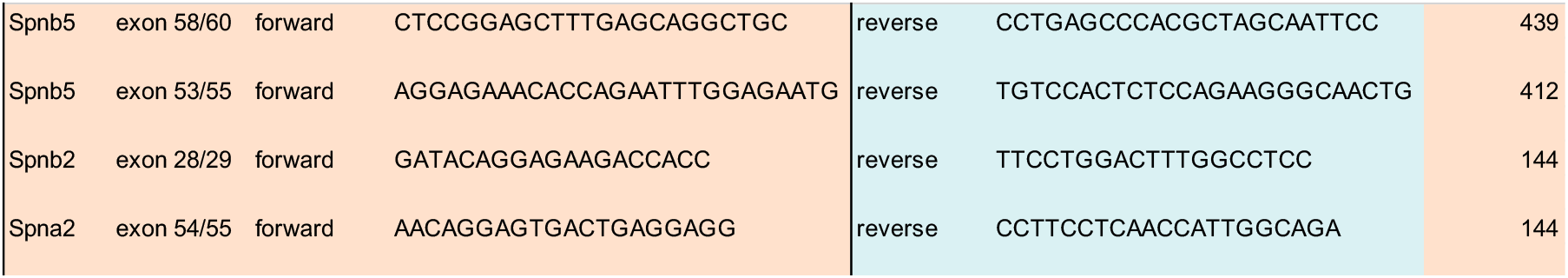
Primers for rt-PCR in WT and Spnb5^(-/-)^ tissue. These primer sets were tested on cDNA from testis isolated from WT and Spnb5^(-/-)^ 3-month-old mice. Because the C-terminus of these large genes tends to be more abundant than N-terminus exons, we designed primer sets that will amplify exons near the poly-A tail.

**Figure S2.**
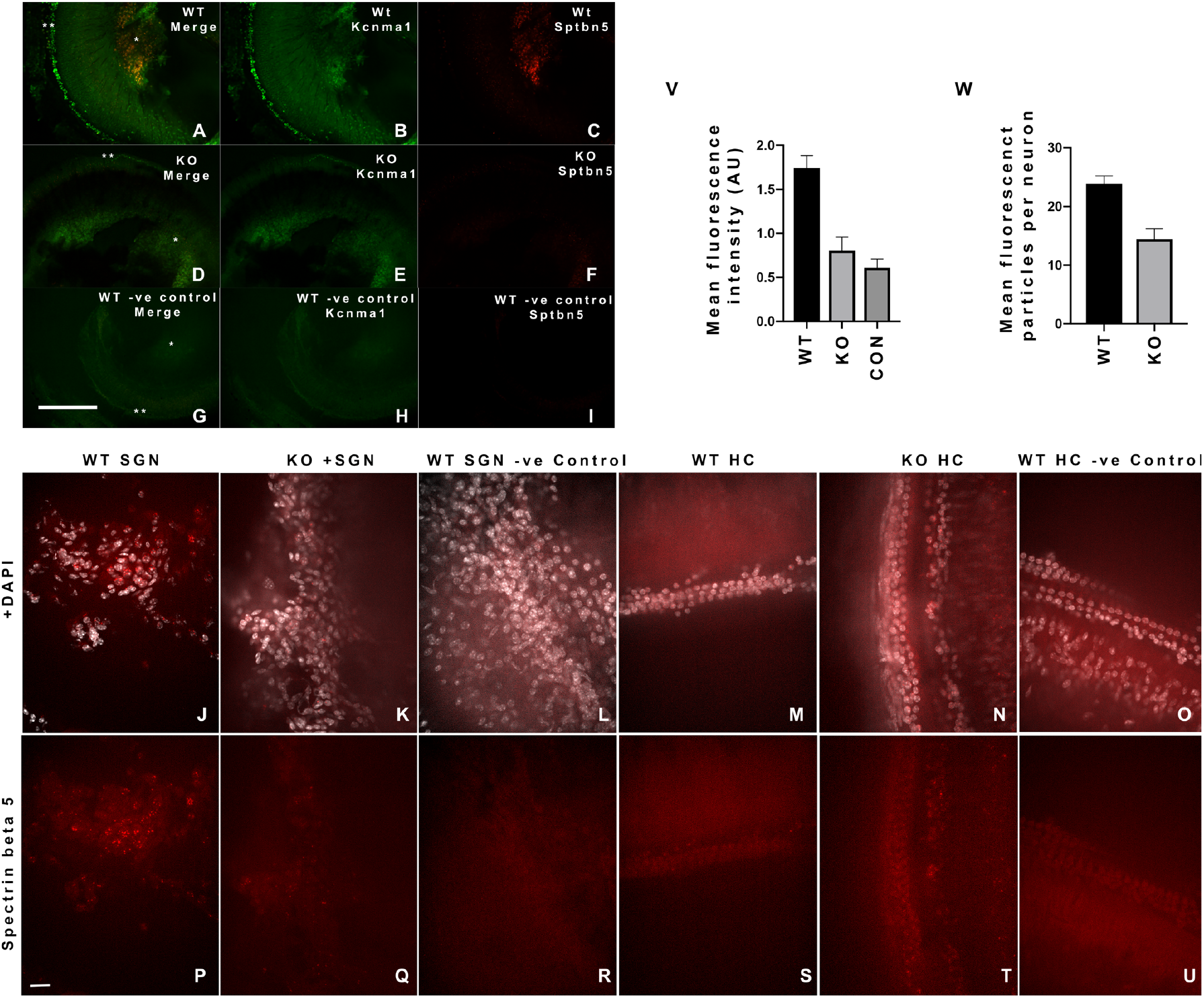
*In situ* hybridization (RNAscope) shows the expression of Spnb5 and Kcnma1 in WT and Spnb5^(-/-)^ cochlear tissue. In the top image, panels (A-F) are a low magnification (10x) comparison of Spnb5 mRNA (red, C & F) and Kcnma1 mRNA (green, B & E) between WT and KO from the apical turn of the organ of Corti. Merged images are displayed in panels (A & D) and the DapB negative control images (-ve control) are shown in panels (G-I). Single asterisks indicate spiral ganglion neurons (SGNs) and double asterisks highlight regions with hair cells (HC). Mean SGN Spnb5 fluorescence intensity of WT animals were significantly different from KO animals and negative controls (V) (WT 1.744 ± 0.13 SEM, n = 9, KO 0.80 ± 0.15 SEM, n = 7, -ve control 0.60 ± 0.09 SEM, n = 5 one-way ANOVA, p<0.001). Areas of SGNs were demarcated based on staining of nuclei (DAPI). Scale bar, 200 μm. The bottom image (J-U) is a single-color, high magnification (40x) for WT and KO tissue. mRNA for Spnb5 was assessed in the SGNs (P-R) and HCs (S-U). The upper panels (J-O) are the same images with DAPI demarcating cell nuclei. Panels (J,P,K,Q) representing Spnb5 mRNA levels (red signal) in WT and KO animals show reduced expression of Spnb5 mRNA in KO animals. Negative control (DapB) images are shown in panels L & R. Of note, Spnb5 is not expressed in all SGNs. In KO animals, the number of Spnb5 granules per neuron, the overall fluorescence intensity of these granules, and the number of neurons with granules were all reduced compared to WT animals. Mean number of Spnb5 mRNA granules per SGN in WT and KO mice (W) were significantly different (WT 23.8 ± 1.3 SEM, n = 23 and KO 14.4 ± 1.7 SEM, n = 28 neurons, respectively, students t test, P <0.001). Here, only those neurons with Spnb5 signal were included. Panels (M,S,N,T) of Spnb5 in HCs of WT and KO cochlea show no signal above DapB background (O & U). Scale bar, 20 μm.

**Figure S3.**
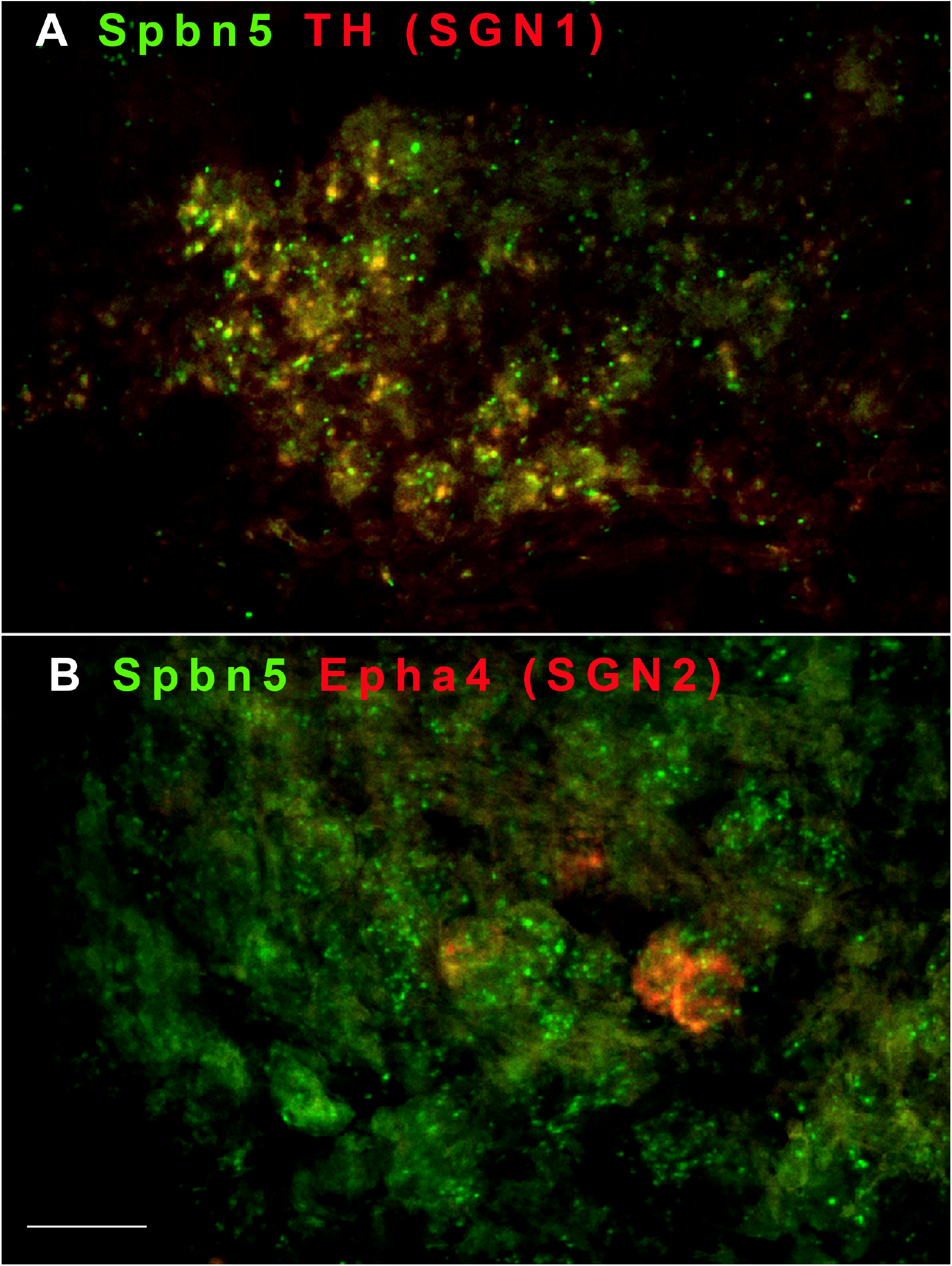
*In situ* hybridization (RNAscope) shows the expression of Spnb5 in SGNs I and II. In the top image, panel (A) is a merged high magnification (40x) comparison of Spnb5 (green) and tyrosine hydroxylase (TH) (red) mRNA. Yellow granules in this image demarcate coincident RNA labeling in SGN Type I cells. Panel (B) is a merged image of Spnb5 (green) and Epha4 (red) signals. Yellow granules are coincident labeling of Spnb5 and Epha4 mRNA in three SGN Type II cells. Scale bar, 20 μm.

### Auditory thresholds and OHC function are similar between Spnb5^(-/-)^ and control mice

To investigate how the loss of Spnb5 impacts auditory information processing, we compared the *in vivo* ABRs of 2-month-old Spnb5^(+/+)^ and Spnb5^(-/-)^ mice. ABR thresholds remained unaltered in Spnb5^(-/-)^ mice across all sound frequencies tested (**Figure 3A**). We also found that distortion product otoacoustic emissions (DPOAEs), a measure of the robustness of OHC function, were unaffected by the loss of Spnb5 (**Figure 3B**). These findings with preserved DPOAE thresholds also extended to suprathreshold responses at sound intensities of 80-90 dB SPL (**Figure 3C**). Since spectrins have been implicated in membrane integrity and protein shuttling, we also assayed prestin expression in the lateral wall using its gating charge movement (NLC) at postnatal day 13 (when prestin expression increases in OHCs) and at two months (Abe et al., 2007). At both these time points, we found NLC to be unaffected in Spnb5^(-/-)^ mice (**Figure 4A**). Our measures of the different parameters of NLC, Q_sp_, V_h_, and z, were similar between WT and Spnb5^(-/-)^ mice (**Figure 4, A-D**).

**Figure 3.**
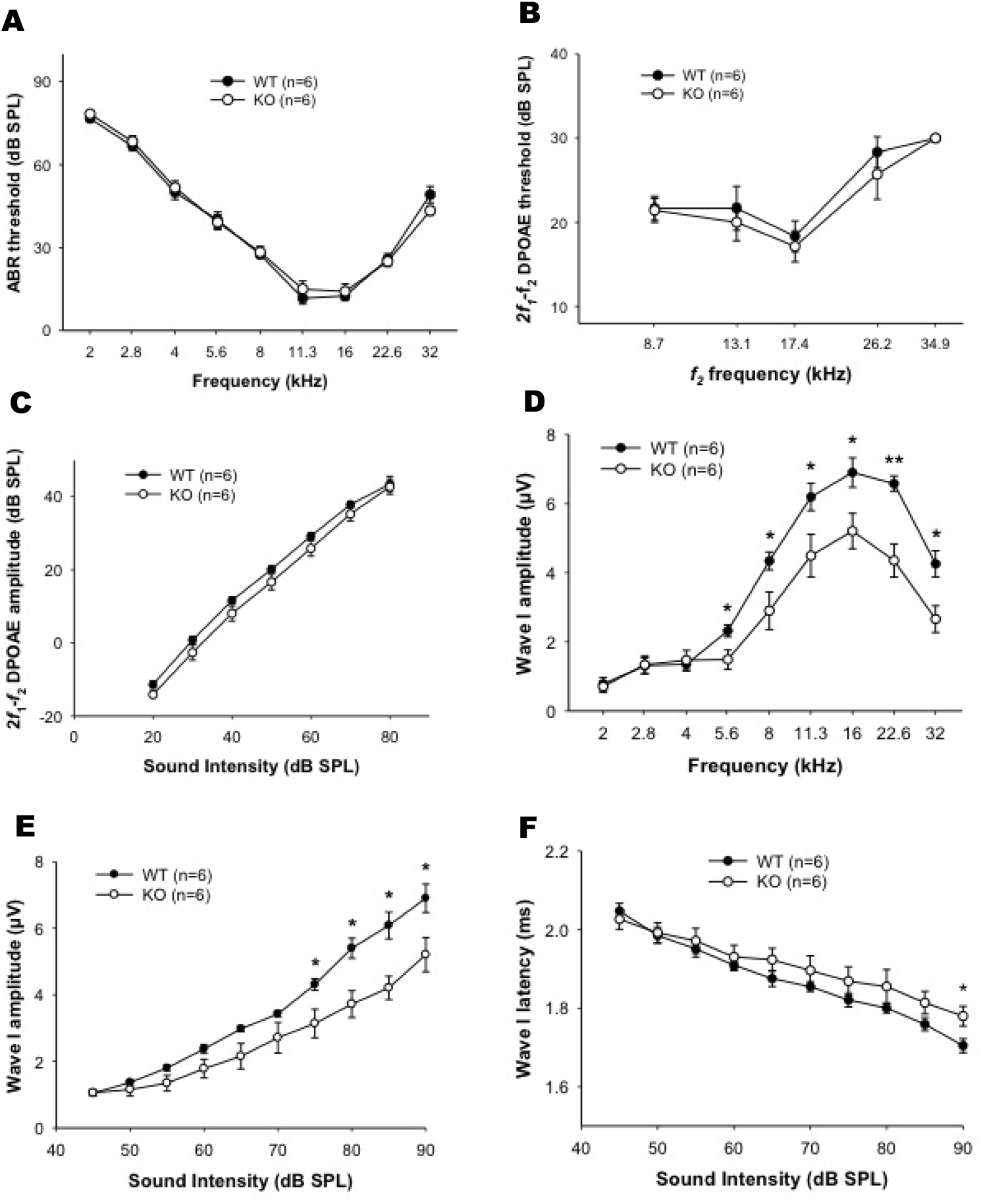
ABR analysis suggests a neuronal and not OHC dysfunction in Spnb5^(-/-)^ mice. **(A) ABR thresholds at frequencies between 2-32 kHz remain unchanged in Spnb5^(-/-)^ mice.** ABR thresholds in Spnb5^(-/-)^ (open circle) and littermate WT (solid circle) mice at 2 months of age are plotted as a function of stimulus frequency. There are no significant differences between WT and Spnb5^(-/-)^ mice at all frequencies tested from 2 to 32 kHz. **(B) Similar 2f_1_-f_2_ DPOAE thresholds are observed in 2-month-old WT and Spnb5^(-/-)^ mice at all f_2_ frequencies tested. (C) 2f_1_-f_2_ DPOAE amplitudes are unaffected in Spnb5^(-/-)^ mice.** Input/output (I/O) function analysis of **2f_1_-f_2_** DPOAE responses in WT (solid circle) and Spnb5^(-/-)^ (open circle) mice as a function of sound intensity at the f2 frequency of 17.4 kHz at 2 months of age are graphed. DPOAE amplitudes in Spnb5^(-/-)^ mice show no significant differences across all sound intensities compared to WT mice. **(D) ABR wave I amplitudes are significantly altered between Spnb5^(-/-)^ and WT at high stimulus intensities at frequencies between 5.6-32 kHz, but not at frequencies between 2-4 kHz.** Shown are wave I amplitudes at 90 dB SPL. There is a consistent reduction in wave I amplitudes at these suprathreshold intensities from 5.6-32 kHz. **(E) ABR wave I amplitudes at 16 kHz are significantly reduced in Spnb5^(-/-)^ mice at high sound intensities (75-90 dB SPL). (F) ABR wave I latencies at 16 kHz are significantly delayed in Spnb5^(-/-)^ mice at 90 dB SPL.** Latencies at all other frequencies are not significantly delayed (data not shown). Data are means +/- SEM. The number (n) of animals per experiment is shown in each graph. Asterisks indicate p<0.05.

**Figure 4.**
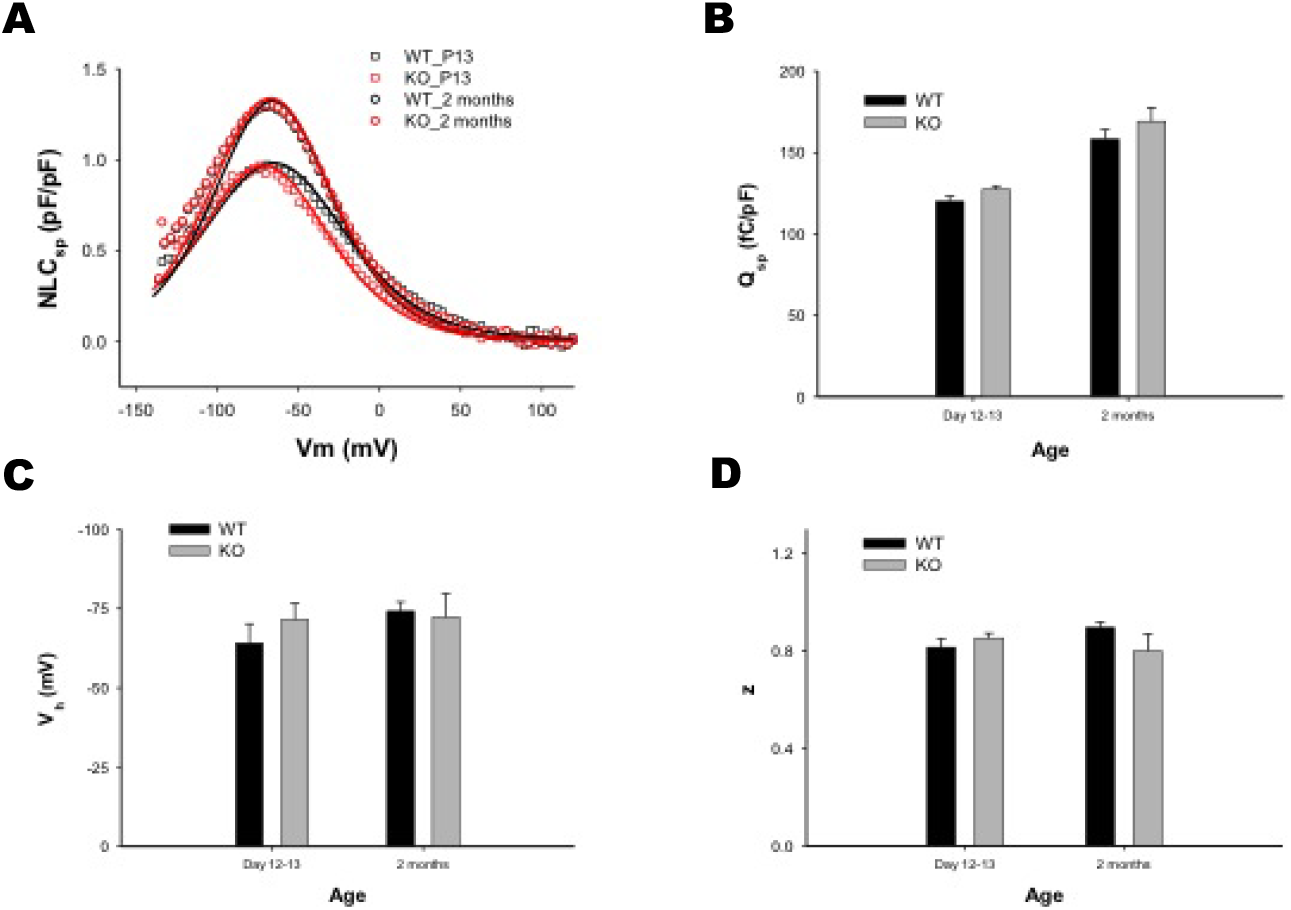
Spnb5^(-/-)^ mice have similar nonlinear capacitance characteristics as controls at two age groups. NLC was measured at postnatal day 13, a midpoint of NLC development, and at two months when hearing is mature. **(A) There are no significant effects of OHC NLC in Spnb5^(-/-)^ mice.** Shown are representative traces of NLC measurement in Spnb5^(-/-)^ and WT mice. There are no differences in NLC between Spnb5^(-/-)^ and WT mice at these two time points. **B, C, and D. NLC parameters are unaffected in Spnb5^(-/-)^ mice.** Shown are means (+/- SEM) of NLC parameters in Spnb5^(+/+)^ and Spnb5^(-/-)^ mice at 13 days and two months, with no significant differences (P>0.05) in all of these measures (Q_sp_, V_h_ and z). **(B) Q_sp_, a specific measure of net charge movement per unit of membrane corrected for linear capacitance. (C) V_h_, a measure of the voltage dependence of charge movement. (D) z, a measure of the slope in NLC voltage dependence and indicative of the unitary charge movement.**

### Loss of βV spectrin affects hearing sensitivity in a manner suggesting a neural processing disorder

The ABR is characterized by a series of electrical waves representing the progressive transfer of the auditory signal from the periphery to the CNS (Song, McGee, & Walsh, 2006). Wave I represents the summated response from the spiral ganglion and auditory nerve (Melcher et al., 1996). While we see no differences in ABR thresholds, we note that wave I amplitudes from 5.6 kHz and upwards are significantly altered between Spnb5^(-/-)^ and WT mice at high stimulus intensities (suprathreshold) (**Figure 3D**). We also analyzed the latency of wave I to investigate any potential changes in the rate of propagation of neural transmission. Wave I latency at 16 kHz was progressively delayed at high sound intensities reaching significance at the highest sound intensity (90 dB SPL) (**Figure 3F**). Latencies at other frequencies were not significantly different from WT.

### Spnb5 plays a critical role in sensorineural processing

What could be the cause of the differences in amplitudes of wave I between WT and Spnb5^(-/-)^ mice? We sought to exclude synaptopathy as a cause of the decrease in wave I amplitudes. For instance, spectrins have been shown to be important in targeting structural elements of the synaptic apparatus as well as maintaining the integrity and compartmentalization of axons and dendrites ((Hammarlund, Jorgensen, & Bastiani, 2007), (Stankewich et al., 2010), (Efimova et al., 2017; Stankewich et al., 2011), (Lorenzo et al., 2019). We compared the number of IHC ribbon synapses in the basal, middle, and apical turn of the organ of Corti between Spnb5^(-/-)^ and WT mice. We used antibody labeling of the marker protein CtBP2 (Bramhall, McMillan, Kujawa, & Konrad-Martin, 2018), a major component of the ribbon synapses, and a well-defined marker of synaptopathy (Sergeyenko, Lall, Liberman, & Kujawa, 2013). We found no differences in CtBP2-labeled synaptic boutons of IHCs in the three cochlear turns between WT and Spnb5^(-/-)^ mice (**Figure 5A**). We then sought to identify morphological differences in afferent nerve numbers as a possible cause of the reduced wave I amplitudes. We found significant reductions in neurofilament labeled, nonmyelinated, afferent fibers in the habenular in all three turns as well as efferent fibers in the basal and middle turns of the cochlea (**Figure 5H, O**).

**Figure 5.**
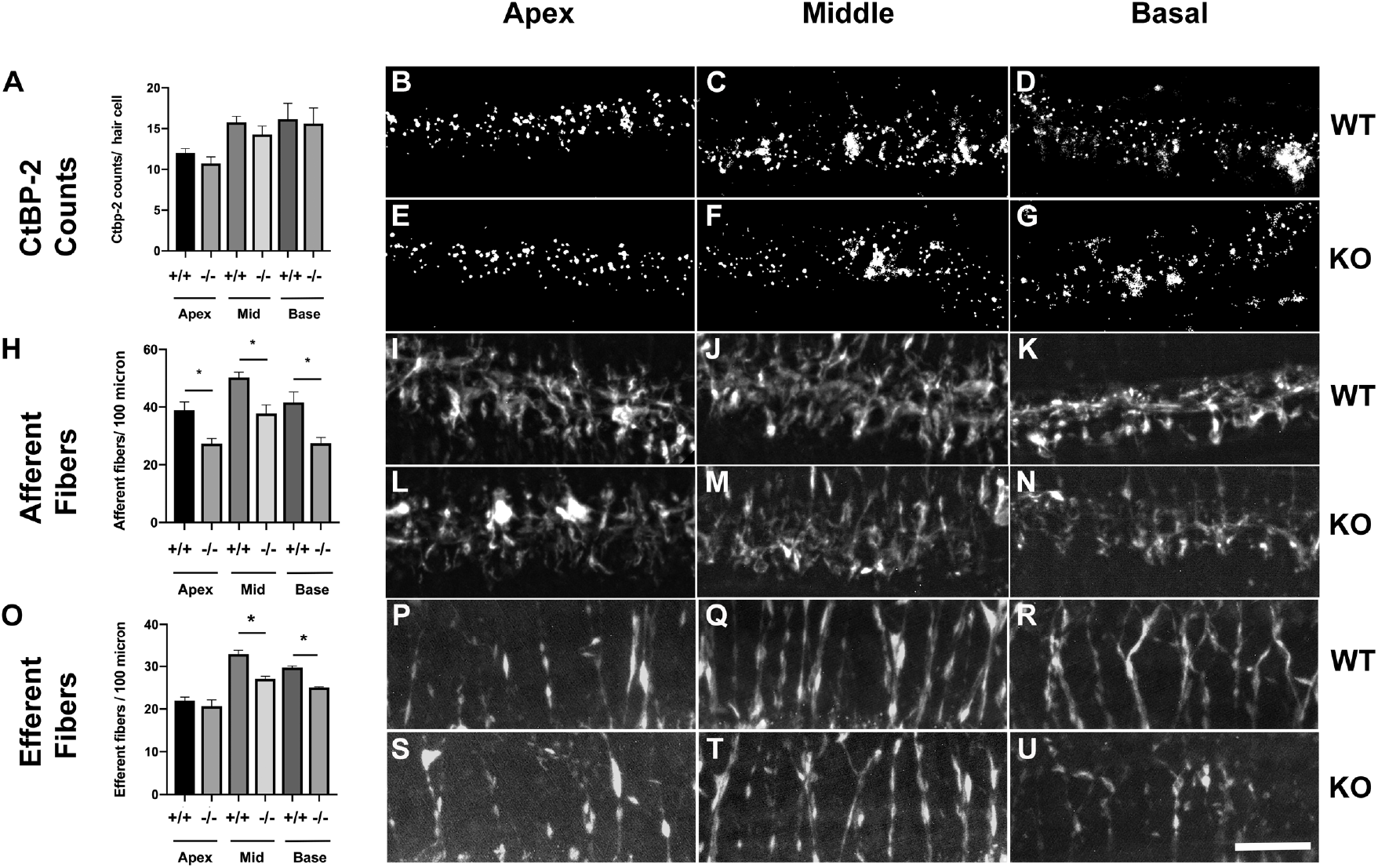
Spnb5^(-/-)^ mice show evidence of neuropathy and no evidence of a synaptopathy. Shown are representative z axis projections of confocal micrographs of WT and βV spectrin KO mice in which synaptic boutons are labeled with antibodies to CtBP2 (**B-G**). **Afferent (I-N) and efferent (P-U) nerve fibers are labeled with antibodies to neurofilament.** The scale bar equates to 20 microns. There is a significant reduction in the density of afferent fibers at the habenular in all three turns and efferent fibers in the cochlear canal of Spnb5^(-/-)^ mice. In contrast, there are no differences in CtBP2-positive bouton counts in all three turns of the cochlea. (**A, H, O) Shown are mean (+/- SEM) counts of synapses per IHC (A), afferent fibers (H), and efferent fibers (O) in all three cochlear turns.** The differences in afferent and efferent counts are significant (* P<0.05, n = 4 cochlea) between WT and Spnb5^(-/-)^ mice in all three turns of the cochlea (afferent), and in the middle and basal turns of the cochlea (efferent). CtBP2 counts between WT and Spnb5^(-/-)^ mice are not significantly different (P>0.05).

## DISCUSSION

In this paper, we elucidate the function of βV spectrin in hearing using a mouse gene deletion model. Removal of exon 2 containing the initiator methionine and an N-terminal 37 amino acids, which, in part, compose its actin-binding domain, results in a mouse with normal ABR thresholds, and normal measures of OHC function including DPOAEs and NLC. Typical DPOAEs are a strong indication of normal OHC electromotility (Shera & Guinan, 1999). The presence of normal NLC measures further reiterates this interpretation. Moreover, we were unable to detect Spnb5 mRNA in the organ of Corti or OHCs.

In contrast, we find a reduction in ABR wave I amplitudes that are concordant with our finding of reduced numbers of nerve fibers. These data point to βV spectrin as critical for the maintenance and function of afferent and efferent nerve fibers. The normal ABR thresholds in the presence of reduced wave I amplitudes we observe in mice have been seen previously in mutations of other proteins and, in particular, in the recovery phase of noise-induced hearing loss (Maison, Pyott, Meredith, & Liberman, 2013) (Kujawa & Liberman, 2009).

How do we explain the preserved ABR thresholds in the presence of reduced wave I amplitudes? Loss of neurons of up to 80% has been shown to be associated with normal thresholds as long as hair cells were functional (Schuknecht & Woellner, 1953). Although speculative, other authors have reasoned that a small increase in sound pressure of 5 dB results in wider recruitment of neurons adjacent to the index sound frequency that could compensate for a 50% loss in afferent neurons using insensitive measures such as compound action potentials and ABRs (Taberner and Liberman 2005), (Lin, Furman, Kujawa, & Liberman, 2011). An important distinction in our data is that, unlike in noise-induced hearing loss models, we see no differences in CtBP2 counts (Liberman & Kujawa, 2017). Our CtBP2 counts and preliminary analysis using nerve counts suggest that the functional effects in these mice lie in the nerve. Ongoing work in our lab includes detailed analysis of high and low threshold fiber counts in these animals, together with studies that will better determine the exact molecular mechanisms which underlie the reduction in dendritic and axonal numbers.

We find absent effects on OHC function in Spnb5^(-/-)^ mice that is consistent with the undetectable expression of these transcripts in the cochlea and OHC in particular. We note similar low/absent Spnb5 RNA expression upon interrogation of both OHC single-cell RNA and OHC RNA expression datasets of adult mice that have been published (Li et al., 2018) and gEAR portal (Shen, Scheffer, Kwan, & Corey, 2015), (Liu, Chen et al. 2018), (Hoa et al., 2020). Together, these data question the validity of βV spectrin antibody staining of OHCs, the principal reason for ascribing its presence in OHCs (Legendre et al., 2008), (Cortese et al., 2017). These papers led to the belief that βV spectrin had an integral role in the cytoskeleton of hair cells. Our assertion that βV spectrin is not important for OHC function is based, in part, on the absent phenotype in the knockout mice. It is reasonable to ask if the knockout mouse resulted in an actual loss of function in the gene. We feel confident that the deletion of the second exon in Spnb5^(-/-)^ resulted in a silencing of the gene. In addition to the neuronal phenotype we describe herein with the Spnb5^(-/-)^ mice, we were also unable to detect Spnb5 transcript from the testis of knockout mice that may relate to degradation of nonsense transcripts and a phenomenon observed in deletion of other genes (Baker & Parker, 2004). RNAscope *in situ* hyridization data also show reduction in the amount of detected mRNA in the SGN of KO animals. While the phenotype of mice is consistent with the absent expression of Spnb5 transcript in OHCs, we have to be mindful that normal OHC function in these mice could be due to other beta spectrins substituting for Spnb5 spectrin. In this context, the presence of a clear auditory phenotype in the Spnb2 spectrin knockout mice would suggest that such an interchange and substitution by other spectrins is, however, not likely. Moreover, Spnb4^(qv/qv)^ spectrin has a neuronal phenotype, similar to Spnb5^(-/-)^.

In conclusion, we present data in this study that βV spectrin plays an important role in auditory function. Our data confirm that βV spectrin is necessary to maintain the integrity or function of afferent and efferent nerves of the peripheral auditory system. Importantly, our data show no evidence that βV spectrin has a function in OHCs, as has been previously suggested.

## MATERIALS AND METHODS

### Mice

All procedures conducted with mice were approved by the Institutional Animal Care and Use Committee at the Yale University School of Medicine. C57BL/6J mice were housed under pathogen-free conditions and maintained as a breeding colony. In our studies, C57BL/6J mice were engineered to harbor a gene knockout of Spnb5 (βV spectrin). The floxed exon 2 of Spnb5 was excised with Cre-recombinase from embryonic stem (ES) cells in culture before injecting them into mouse blastocysts under the guidance of the Yale Gene Targeting Center. This construct was designed to create a constitutive loss of βV spectrin. For genotyping animals, tail snips were collected from mice at their weaning age. Genomic DNA was prepared by an overnight 56°C incubation in tail lysis buffer 50mM Tris pH 8.0, 50mM KCL, 2.5mM EDTA, 0.4% Igepal, 0.4% Tween 20 supplemented with Proteinase K (400 μg/ml). gDNA was assayed by PCR using the following primer set: Forward 5’-CTGTCCGCCAAAGAGCAGGGGCCATG-3’ Reverse 5’-GATGTGGGGTCAGCCCAGCATCTCC-3’ and PCR amplimers were visualized on a 2% agarose gel.

### OHC isolation

Prestin-YFP mice of 2-3 months of age were euthanized, the cochleae extracted, and the organ of Corti dissected in ice cold 1x DPBS. OHCs were dissociated by enzymatic digestion with pre-warmed 0.05% trypsin for 8 minutes at 37°C followed by gentle trituration in 10% FBS in DMEM with a 1 mL pipette tip. The cell suspension was spun at 1500 g for 10 minutes. The cell pellet was resuspended in DMEM supplemented with 10% FBS and then filtered through a 40-micron cell strainer. The cells were FACS sorted, and YFP-positive OHCs were collected.

### mRNA detection

Total RNA from mouse testis and the whole brain was either obtained from Takara or prepared from 100 mg of testis tissue from 3-month old WT and βV KO C57BL/6J animals with Trizol reagent (Chomczynski & Mackey, 1995). cDNA was prepared with Superscript IV reverse transcriptase (ThermoFisher Scientific, Waltham, MA,) and quantified with a Nanodrop spectrophotometer. Standard procedures followed reagent manufacturer’s directions for first-strand synthesis. The organ of Corti and OHCs (prepared as described above) were isolated and proteolyzed with 1 unit of proteinase K. After inactivating at 95°C for 5 minutes, non-purified RNA was treated with Superscript IV to generate cDNA. Gene-specific PCR primers bridging consecutive exons were designed to ensure the exclusion of genomic DNA from amplification in **Table 1**.

### RNAscope *in situ* Hybridization

RNAscope *in situ* hybridization was performed on cochlear whole mount tissue using RNAscope Multiplex Fluorescent v2 Assay (Advanced Cell Diagnostics (ACD), Inc. Newark, CA, USA) according to the manufacturer’s instructions (Document # 323100-USM). RNAscope Probes and RNAscope Mulitplex Fluorescent Reagent Kit v2 (Catalog # 323100) required for the assay were purchased directly from ACD. Briefly, the cochlear apical turn of the organ of Corti was dissected in 1x DPBS and fixed in 4% PFA. Cochlear tissue was pre-treated with hydrogen peroxidase for 10 min at room temperature (RT), then with Protease Plus for 15 min at 40°C in the HybEZ II Oven (ACD). Tissues were washed thrice with distilled water for 5 min each at RT. The RNAscope Assay used the following probes: target probe for Spnb5 (Catalog # 548431); positive control probe, Kcnma1 (Catalog #476251-C2); and negative control probe, DapB (Catalog #310043). For SGNs the two probes used were Epha4 (Catalog #419081) and tyrosine hydroxylase (Catalog #317612-C3). All probes were pre-warmed (10 min at 40°C), brought to RT, then hybridized to the tissue with an 8 h incubation at 40°C. After the tissue was stored in 5x saline sodium citrate (SCC) overnight at RT, hybridized probe signal was amplified at 40°C with the consecutive addition of three amplifiers: AMP1 (30 min), AMP2 (30 min), and AMP3 (15 min). HRP signal was developed with a 15 min incubation in HRP-C1 at 40°C. For a single label experiment, the TSA plus fluorophore CY3, diluted (1:50), (TSA Fluorescence Kit, Catalog # NEL760001KT; PerkinElmer, Inc., MA, USA) was added for 30 min at 40°C, followed by a 15 min treatment with HRP blocker at 40°C. For double fluorescent experiments, probes were developed using both HRP-C1 and HRP-C2 with alternating treatments of each fluorophore (fluorescein and CY3) and the HRP blocker step. Between steps, tissue was washed twice with 1x Wash Buffer for 2 min each at RT. Finally, tissue was mounted with Prolong Gold antifade reagent with DAPI (Thermo Fisher Scientific, OR, USA) on a glass bottom dish, and fluorescent signal was visualized and imaged using a confocal microscope (Zeiss Observer Z1).

### Auditory brainstem responses (ABR)

ABR measurements were conducted in a manner similar to that described by Song *et. al*. (El-Hassar et al., 2019). Briefly, animals were anesthetized with 480 mg/kg, i.p., chloral hydrate, and all recordings were conducted in a sound-attenuating chamber (Industrial Acoustics Corp., Bronx, NY). A customized TDT System 3 (Tucker-Davis Technologies Inc., Alachua, FL) was used for ABR recordings. Subdermal needle electrodes (Rochester Electro-Medical, Lutz, FL) were positioned at the vertex (active, noninverting), the right infra-auricular mastoid region (reference, inverting), and the left neck region (ground). Differentially recorded scalp potentials were bandpass filtered between 0.05 and 3 kHz over a 15 ms epoch. A total of 400 responses were averaged for each waveform for each stimulus condition.

Symmetrically shaped tone bursts were 3 ms long (1 ms raised cosine on/off ramps and 1 ms plateau). All acoustic stimuli were delivered free field via a speaker (EC1 Electrostatic Speaker, Tucker-Davis Technologies) positioned 10 cm from the vertex. Stimulus levels were calibrated using a 0.5-inch condenser microphone (model 4016, ACO Pacific, Belmont, CA) positioned at the approximate location of the animal’s head during recording sessions and are reported in decibels of sound pressure level (SPL; referenced to 20 μPa). Stimuli of alternating polarity were delivered at a rate of 21/s. Tone burst responses were collected in half-octave steps ranging from 32 to 2.0 kHz. Stimuli were presented in 5 dB decrements of sound intensity from a maximum level of 90 dB SPL. The ABR threshold was defined as the lowest intensity of sound level capable of evoking a reproducible, visually detectable response. Latencies and amplitudes of the initial ABR wave were determined at a frequency of 16 kHz, one of the most sensitive frequency of hearing in mice. Latencies and amplitudes of wave I at 90 dB SPL were also measured at all the other frequencies. The analysis was performed in BioSigRP, a digital signal processing software (TDT), on traces with discernible peaks by setting cursors at the maxima and minima of the peaks. Latency was defined as the time from the onset of the stimulus to the peak, while amplitude was measured by taking the mean of the ΔV of the positive and negative deflections of the ABR wave.

### Distortion product otoacoustic emissions (DPOAEs)

The acoustic stimuli for DPOAEs were produced, and responses were recorded using a customized TDT System 3 (Tucker-Davis Technologies Inc., Alachua, FL) controlled by BioSigRP (TDT). To deliver acoustic stimuli and record the responses, two speakers (MF1 Multi-Field Magnetic Speakers, TDT) and a microphone probe (ER-10B+, Etymotic Research, Elk Grove Village, IL) were connected by short PVC tubing, and the microphone probe was inserted into the external auditory meatus of the anaesthetized mouse via a tapered plastic pipette tip. Two simultaneous continuous pure tones (*f*_1_ and *f*_2_) that have a frequency ratio of 1.2 (*f*_2_/*f*_1_) and equal sound level (*L*_1_ = *L*_2_) were delivered at center frequencies of 8, 12, 16, 24 and 32 kHz with sound levels from 80 to 20 dB SPL in 10 dB decrements. Stimuli duration was 83.88 ms at a repetition rate of 11.92 per second. The acquired DPOAE responses were averaged 100 times. The DPOAE of interest was at 2*f*_1_-*f*_2_, the largest and most prominent DPOAE. The DPOAE threshold was defined as the lowest sound intensity capable of evoking a visually detectable 2*f*_1_-*f*_2_ signal above the noise floor. DPOAE amplitudes were analyzed offline in BioSigRP by setting cursors at the peak of the 2*f*_1_-*f*_2_ signals.

### Nonlinear capacitance (NLC)

Nonlinear capacitance was conducted in a fashion much like that previously described (Song & Santos-Sacchi, 2016). C57BL/6J WT and βV spectrin KO littermates at postnatal day 13 and at 2 months of age were compared. Apical turns of the organ of Corti were isolated from each and recorded with ionic current blocking solutions to minimize interference on measures of membrane capacitance. The extracellular solution contained (in mM): 100 NaCl, 20 tetraethylammonium (TEA)-Cl, 20 CsCl, 2 CoCl_2_, 1 MgCl_2_, 1 CaCl_2_, 10 HEPES, pH 7.2. The intracellular solution contained (in mM): 140 CsCl, 2 MgCl_2_, 10 HEPES, and 10 EGTA, pH 7.2. Pipettes had resistances of 3-5 MΩ. Gigohm seals were made, and stray capacitance was balanced out with amplifier circuitry prior to establishing whole-cell conditions. A Nikon Eclipse E600-FN microscope with 40× water immersion objective lens was used to observe cells during voltage clamp. Whole-cell voltage-clamp recordings were performed from all three rows of OHCs on whole-mount organ of Corti. OHCs were recorded at room temperature using jClamp software and an Axopatch 200B amplifier (Molecular Devices, LCC, San Jose, CA). Data were low pass filtered at 10 kHz and digitized at 100 kHz with a Digidata 1320A (Molecular Devices, LCC, San Jose, CA). In order to extract Boltzmann parameters, capacitance-voltage data were fit to the first derivative of a two-state Boltzmann function.

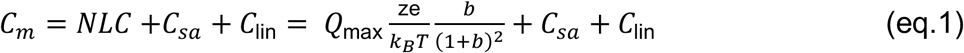

where 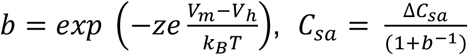

Q_max_ is the maximum nonlinear charge moved, V_h_ is the voltage at peak capacitance or equivalently, at half-maximum charge transfer, V_m_ is R_s_-corrected membrane potential, z is valence, C_lin_ is linear membrane capacitance, e is the electron charge, kB is Boltzmann’s constant, and T is the absolute temperature.

### Immunohistochemistry

For immunohistochemistry, staining was performed on fixed and decalcified three turns of the organ of Corti to compare the number of neuronal fibers and synapses between all three cochlear turns in KO and WT mice. Temporal bones were fixed in 4% PFA overnight at 4°C and then decalcified in 120 mM ethylenediaminetetraacetic acid (EDTA) for 2 days. Whole-mount of the organ of Corti was dissected, washed in PBS 0.05% Tween 20, 0.05% Triton-X 100 (incubation buffer) three times for 3 minutes each, and incubated with primary antibody in incubation buffer overnight at 4°C with mouse anti-CtBP2 antibody (1:50, BD Transduction, San Jose, CA, cat# 612044), chicken anti-neurofilament (NF) antibody (1:500, MilliporeSigma, Billerica, MA, cat# AB5539). After three washes in incubation buffer, the tissue was incubated with secondary antibody in incubation buffer for 1 hour at room temperature. The secondary antibodies used included donkey anti-rabbit Alexa Fluor 568/647 (1:200; Thermo Fisher Scientific, cat # A-10042/A-31573), goat anti-mouse Alexa Fluor 568/647 (1:200; Thermo Fisher Scientific, cat # A-21124/A-21235) and goat anti-chicken Alexa Fluor 488/594 (1:200; Thermo Fisher Scientific, cat # A-11039/A-11042). After antibody treatment, cells were washed in incubation buffer and mounted in Vectashield (Vector Laboratories, Inc., Burlingame, CA), and viewed using a Zeiss confocal laser-scanning microscope (Oberkochen, Germany).

Nerve (NF positive) and synaptic vesicle (CtBP2 positive) counts were performed posthoc in Fiji (ImageJ). Briefly, z stacks of images were separated by probes, background-subtracted (using a 50-pixel rolling ball radius), and z sections containing the images of interest identified for further analysis (slice keeper). CtBP2 labeled IHCs and NF labeled afferent fibers surrounding IHCs in proximity to the habenula and efferent fibers in the cochlear canal were counted. Z projections (sum) of the respective sections were obtained and automated thresholds determined (max entropy). Counts of particles (synaptic vesicles) immunopositive for CtBP2 larger than 0.1 microns were scored. Neuronal numbers were established by counting the peaks in a profile plotted perpendicular to neural fibers in the cochlea canal. For afferent nerve counts the profile was plotted at the midpoint of nerve densities.

### Statistical Analysis

Data were analyzed using FIJI, Microsoft Excel, SigmaPlot and GraphPad Prism. The measures of variability are depicted as error bars +/- the SEM and stated numerically in the figure legends. Unpaired, one or two-tailed Student’s t-test were performed using both Excel and GraphPad Prism to make comparisons between WT and Spnb5^(-/-)^ data points. P-values are indicated in the figure legends to two significant figures. The sample size (n) is stated in the figures.

## AUTHOR CONTRIBUTIONS

MCS, JPB, JSM, DN and JSS conceptualized the project. PRS and JSM generated the Spnb5 deleted mice. MCS and JPB conducted animal breeding, tissue collection and genotyping. MCS, JPB, SK, WT, AS and DN performed gene expression and immunohistochemistry analysis. JPB and LS performed assays to assess hearing function. MCS and DN wrote the manuscript, and all authors contributed to its editing and refinement.

## ACKNOWLEDGEMENTS

These studies were supported by grants R01 DC 008130 and R01 DC 008130 from the National Institute of Health to JSS and DN and grants P01 NS35476 (project 4) and R01 HL28560 to JSM. Other support includes the Raymond Yesner Endowment to Yale University (JSM). We also would like to thank Lan Ji for her technical assistance.

